# EEG-VLM Toolbox: Extending voxel-based lesion mapping to multi-dimensional EEG data

**DOI:** 10.1101/2024.10.25.620269

**Authors:** Richard Hardstone, Lauren Ostrowski, Aliceson N. Dusang, Eduardo López-Larraz, Jessica Jesser, Sydney S. Cash, Steven C. Cramer, Leigh R. Hochberg, Ander Ramos-Murguialday, David J. Lin

## Abstract

Focal brain lesions (such as with stroke) cause functional changes in local and distributed neural systems. While there is a long history of post-stroke neurophysiological assessment using electroencephalography (EEG), the observed neurophysiological changes have rarely been related to specific lesion locations. Therefore, the relationships between anatomical injury and physiological changes after stroke remain unclear. Voxel-based lesion symptom mapping (VLSM) is a tool for statistically relating stroke lesion locations to “symptoms”, but current VLSM methods are restricted to symptoms that can be defined by a single value. Therefore, current VLSM techniques are unable to map the relationships between anatomical injury and multidimensional neurophysiological data such as EEG, which contains rich spatio-temporal information across different channels and frequency bands.

Here we present a novel algorithm, EEG Voxel-based Lesion Mapping (EEG-VLM), that produces the set of significant relationships between precise neuroanatomical injury locations and neurophysiology (defined by a cluster of adjacent EEG channels and frequency bands). Further, the algorithm provides statistical analyses to define the overall significance of each neural structure-function relationship by correcting for multiple comparisons using a permutation test. Applying EEG-VLM to a dataset of recordings from chronic stroke patients performing a cued upper extremity movement task, we found that subjects with lesions in frontal subcortical white matter have reduced ipsilesional parietal cue-evoked EEG responses. These results are consistent with damage to a frontal-parietal network that has been associated with impairments in attention. EEG-VLM is a novel and unbiased method for relating neurophysiologic changes after stroke with neuroanatomic lesions. In the context of focal brain lesions associated with neurological impairments, we propose that this method will enable improved mechanistic understanding, facilitate biomarker development, and guide neurorehabilitation strategies.

## Introduction

Injury to the structure of the brain (i.e., focal neuroanatomical lesions as in stroke) causes complex changes to the functioning of local (Sharp et al. 2000) and distributed neural systems (Carrera and Tononi 2014). Since the introduction of EEG, recordings have been performed to characterize the cortical neurophysiological changes that occur after stroke (Walter 1936; Macdonell et al. 1988; Bentes et al. 2018; Cassidy et al. 2020; Bönstrup et al. 2019; Saes et al. 2020). Before the availability of non-invasive imaging, this research was primarily focused on relating EEG to underlying (post-mortem assessed) lesions, to determine whether EEG could be used to localize lesions (Walter 1936; Williams and Gibbs 1938; Case 1938; van der Drift 1957; Van Der Drift 1961). However, since the ability to perform non-invasive imaging of lesions has become commonly available, the primary focus of post-stroke EEG research has shifted to how metrics derived from neural activity relate to the severity of specific impairments (Bönstrup et al. 2019; Cassidy et al. 2020; Saes et al. 2020) or can predict the extent to which patients will recover (Cuspineda et al. 2007; Bentes et al. 2018; Rogers et al. 2020) or respond to a restorative therapy(Wu et al. 2015). The question of how EEG patterns are related to the precise neuroanatomy of the underlying lesion (Fig. 1A) is now rarely addressed. The few post-stroke EEG studies (Macdonell et al. 1988; Fanciullacci et al. 2017; Shreve et al. 2019; Cassidy et al. 2020) that address lesion characteristics use relatively coarse characterizations of the lesion (e.g., lesion size, cortical lobe, or region of interest), or reduce the multi-dimensional EEG activity to a single value for each patient, which can lead to erroneous conclusions (Fig. 1B).

**Figure 1:**
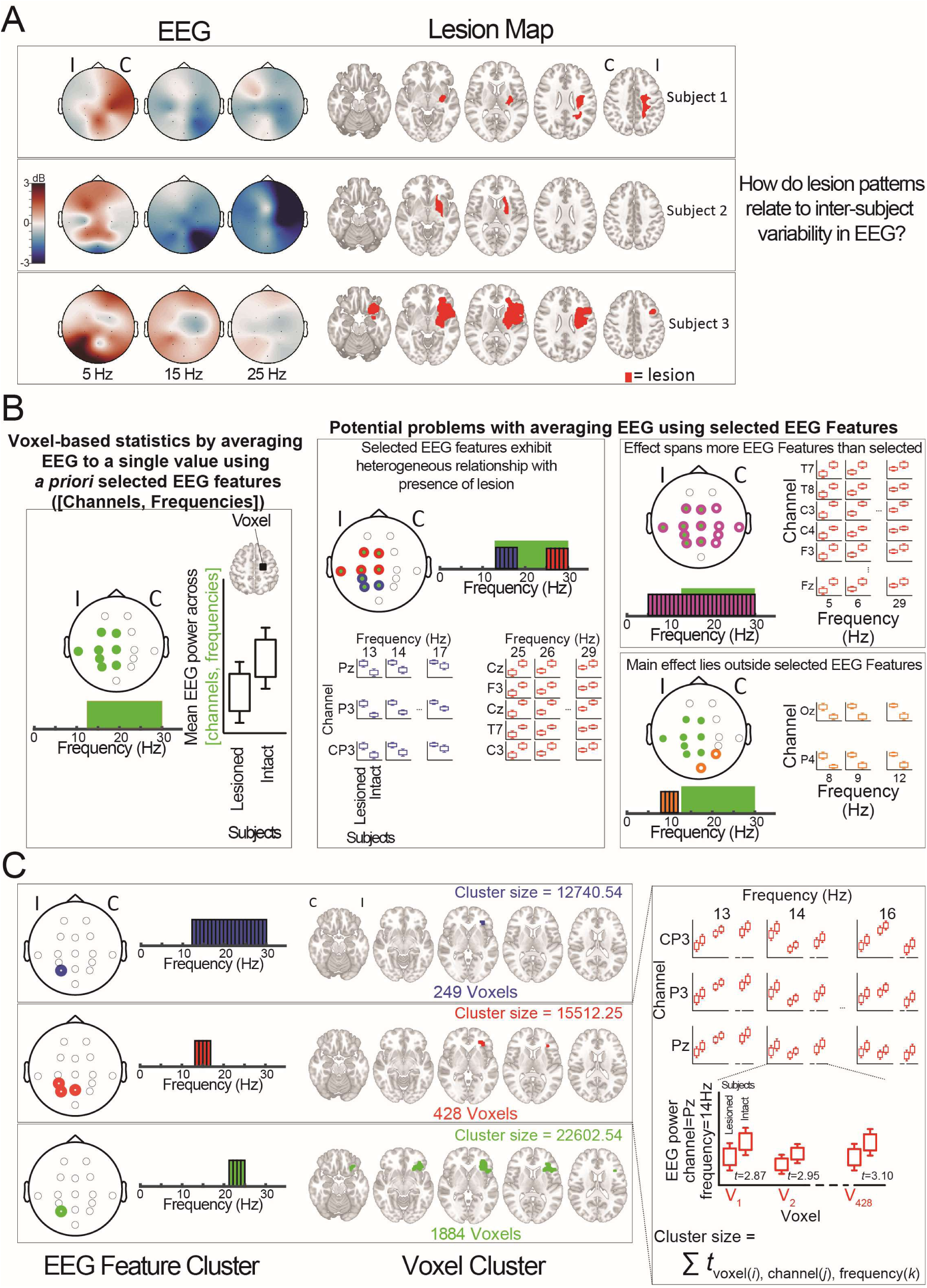
Motivation for linking post-stroke neurophysiology with lesion neuroanatomy. **(A)** Stroke patients exhibit variability in which anatomical regions are lesioned and in their neurophysiology (*Left*) EEG activity exhibits rich and complex spatial and spectral variability across stroke patients (*Right*) Stroke lesions, even those broadly involving same vascular territory and anatomical regions, also show variability across patients, which can be seen at the voxel scale **(B)** (*Left*) To apply current voxel lesion mapping statistics to EEG, requires reducing all EEG Features (power at different channels and frequencies) to a single value. (*Right*) Three different examples where averaging many EEG features (channel, frequency) to a single value loses information, and risks making erroneous interpretations. **(C)** Goal of this paper is to design a method to link post-stroke neurophysiology with neuroanatomy that considers both the rich spatio-temporal information in the EEG and the fine-scale detail of the lesion. The output of this method are interpretable “clusters” of EEG features (e.g. a set of EEG Features ([channels, frequencies]) and lesion voxels where for each voxel, the group of subjects who have that voxel lesioned have significantly different EEG activity (for each EEG feature) to the group who do not have that voxel lesioned. Each [EEG Feature cluster, Voxel cluster] will be associated with a cluster size which can be used to rank effect sizes of different clusters and determine significance. Here three different clusters are shown, with the blue cluster associating EEG activity on one channel across a large frequency range, with presence of a small lesion. With more restricted frequency range (red or green cluster), the association is with presence of a lesion in a larger area, with the red cluster also applying across multiple adjacent EEG channels. (I=Ipsilesional, C=Contralesional).

Since the 19^th^ century, lesion mapping has been a fundamental tool in neuroscience for localizing functions to specific brain regions and applicability and precision have steadily advanced. Early studies analyzed the functional impairments caused by lesions using post-mortem analysis (Broca 1861; Wernicke 1874), ablations in animal models (Leyton and Sherrington 1917), or intra-operative electrical stimulation (Penfield and Rasmussen 1950). However, it was not until the latter half of the 20^th^ Century that a series of methods (pneumoencephalography (Ziegler 1954), isotope mapping (Benson and Patten 1967), Computed Tomography (CT) (Mohr, Watters, and Duncan 1975) and Magnetic Resonance Imaging (MRI) (DeWitt et al. 1984)) were introduced that could non-invasively image lesions which allowed more patients to be studied with progressively finer resolution. To take advantage of the fine-grained imaging resolution of these lesions, new statistical methods (voxel-based lesion symptom mapping, VLSM (Bates et al. 2003) were developed that could map the association between lesions and symptoms at the voxel-level. VLSM allows brain-wide associations to be mapped, moving away from previous *a priori* defined Region-Of-Interest (ROI) based analysis, which has several potential drawbacks including the possibility that the ROI contains multiple subregions that correlated differently with the “symptom”, that the ROI is only a small part of the relevant functional region, or that the relevant functional region is outside the pre-specified ROI. Current VLSM techniques enable brain wide mapping of behavioral metrics, however the behavioral metric must only be a single value per patient. While methods exist that take advantage of the multivariate nature of lesion maps to predict a single behavioral metric (Zhang et al. 2014; Pustina et al. 2018; DeMarco and Turkeltaub 2018; Wiesen, Karnath, and Sperber 2020), there currently exists no method that can take advantage of behaviors or symptoms that are multidimensional.

As a neurophysiological assessment tool, EEG contains rich multidimensional information: over time and across different channels and frequencies (Fig. 1A). To apply current VLSM techniques to this data would require reducing all of this information to a single value, based on *a priori* selection of the channels and frequencies that are of interest. This approach has significant limitations. For example, one might select beta activity (as defined by the classical beta band, 13-30) at electrodes picking up activity from the motor cortex (Fig. 1B, *Left*) to investigate which neuroanatomical regions cause changes across subjects in this motor-related rhythm band (Rossiter, Boudrias, and Ward 2014; Khanna and Carmena 2015; Tang et al. 2020). Applying VLSM in this way can draw incorrect conclusions (Fig. 1B, *Right*), because 1) it assumes that the EEG channels and frequencies selected are homogenous (where there may be subsets of these channels and frequencies showing opposite effects), 2) it assumes the effect only spans the region being studied, but it may be much broader (across more channels and/or frequencies) and so should not be viewed as specifically a motor-related beta effect, and 3) it will not be able to pick up on effects outside the selected region. Therefore, a lesion mapping method that does not require *a priori* selection of EEG features (channels, frequencies) could more accurately capture the changes in neural activity that are caused by particular lesions.

In this paper, we present a new method that discovers statistically significant relationships (Fig. 1C) between multidimensional EEG data (Fig. 1A, *Left*), with 3D spatial information found in lesion maps (Fig. 1A, *Right*). Building up from lesion mapping of zero-dimensional (scalar) EEG data (conceptually equivalent to VLSM, (Fig. 2)) we provide a detailed explanation of the additional steps required to discover relationships that are interpretable for one-dimensional (Fig. 3) and N-dimensional EEG data (Fig. 4). We apply the method to whole-brain lesion maps, and broadband EEG data across the whole cortical surface using a previously recorded dataset of EEG and anatomic lesions from chronic stroke patients during a cued-movement task (Fig. 5). As EEG data during movement might be more contaminated by movement, eye and muscle artefacts(López-Larraz et al. 2018; Bibián et al. 2022), we analyzed the 500 ms post-cue time window to determine the effects of lesions on neural activity. We found a significant relationship between the presence of a lesion in frontal white matter and reduced post-cue evoked responses in an ipsilesional parietal electrode (Fig. 6), consistent with the importance of attentional processes in cued movement tasks. The novel method presented can be used to discover associations between neurophysiology and neuroanatomy, or more generally between multi-dimensional behaviors and neuroimaging, in a hypothesis-free or hypothesis-driven fashion.

**Figure 2).**
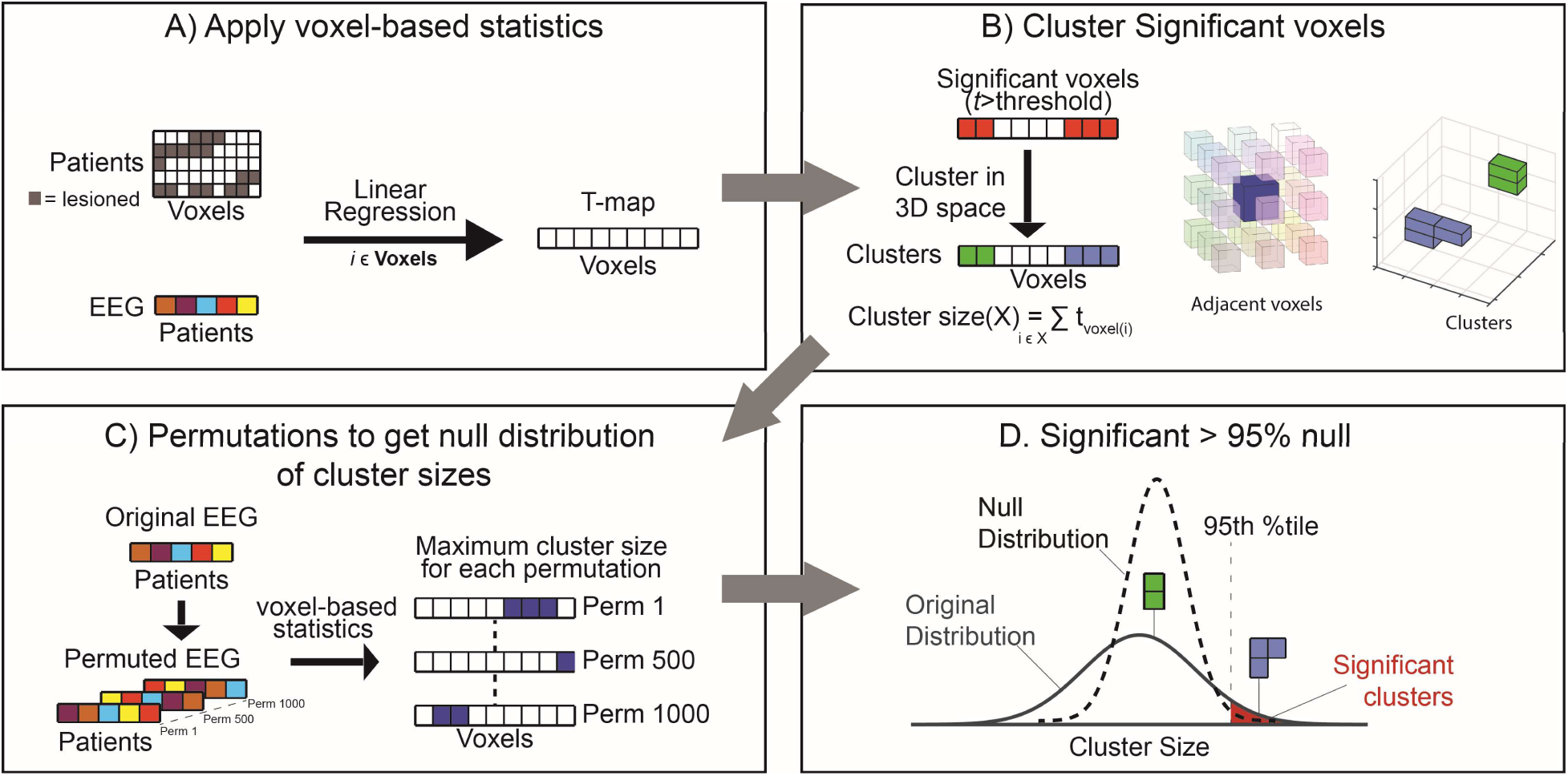
Steps to relate zero-dimension (scalar) EEG data to lesions. **(A)** Input data consists of a set of patients who have a lesion mask (containing all lesioned voxels) and a single value derived from EEG (treated in this case as the “behavior”. For each voxel, linear regression is applied to define the relationship between EEG and lesion presence, resulting in t-statistic, and p-value for each voxel (t-map). **(B)** Significant lesion clusters are calculated by thresholding the t-map (e.g. *p*<0.05) and then finding contiguous voxels in 3D space. The size of each cluster is the sum of the t-statistics from all voxels in the cluster. **(C)** To test the significance of the lesion clusters, a null distribution of lesion cluster sizes is required (i.e. what size of cluster can be expected by chance). To accomplish this a permutation approach is taken, where for each permutation the EEG values are shuffled amongst subjects, and then the same steps (A) and (B) applied to find a set of clusters for each permutation. The maximum cluster size from each permutation forms the null distribution. **(D)** Significant cluster sizes from the unpermuted data are those that are larger than a set proportion of the null distribution (e.g. > 95% of null would correspond to alpha = 0.05).

**Figure 3).**
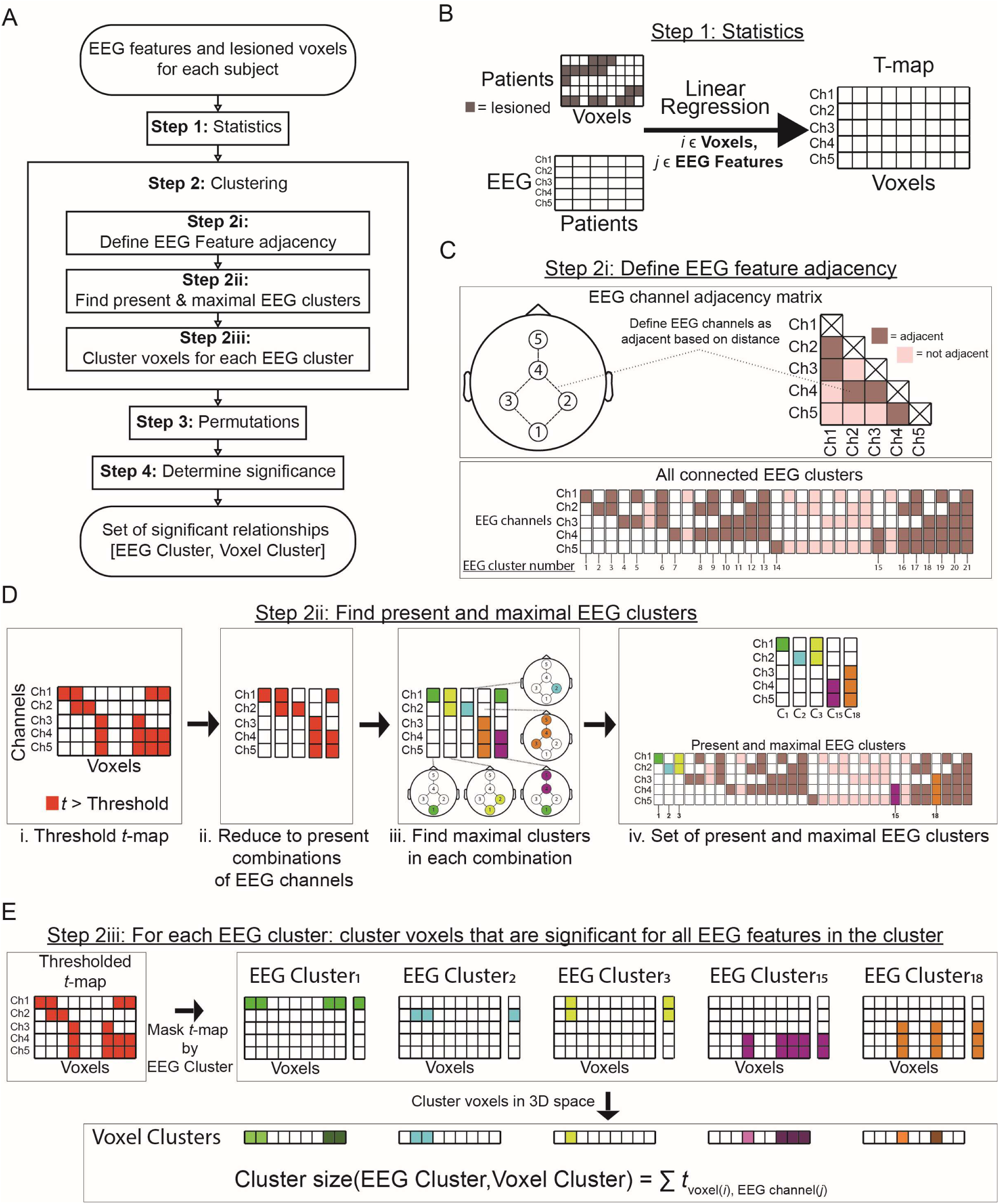
Additional steps to relate 1-dimension EEG data to lesions. **(A)** Additional steps (Steps 2i, 2ii and 2iii) are required to relate lesion patterns to EEG, when EEG consists of more than one value per patient (i.e. > zero dimensions). **(B)** Compared to the 0D case (Fig. 2A), EEG data now consists of a set of EEG features per patient (e.g., one value per EEG channel). Statistics are now applied for each combination of voxel and EEG feature, with resulting t-map and p-values having dimensions of (voxels x #EEG features). **(C)** EEG features are not independent as they show spatial-correlations. Therefore, we can cluster combinations of EEG channels based on spatial distance by defining an adjacency matrix. i.e., EEG channels that are closer to each other will be adjacent. **(D)** Steps to find the set of maximal EEG feature clusters that are present in the thresholded t-map. Only investigating EEG Feature clusters that are present in the thresholded t-map, reduces computation time. The total number of combinations of EEG features will be >= the number of combinations of EEG features which are connected (via adjacency matrix) >= the number of combinations of EEG features that are connected, present and maximal. **(E)** For each maximal EEG cluster present, the thresholded *t*-map is masked to voxels that are significant for all EEG features in that cluster. These voxels are then clustered in 3D space, and the cluster size is the sum of the t-statistics across all [voxel, EEG features].

**Figure 4).**
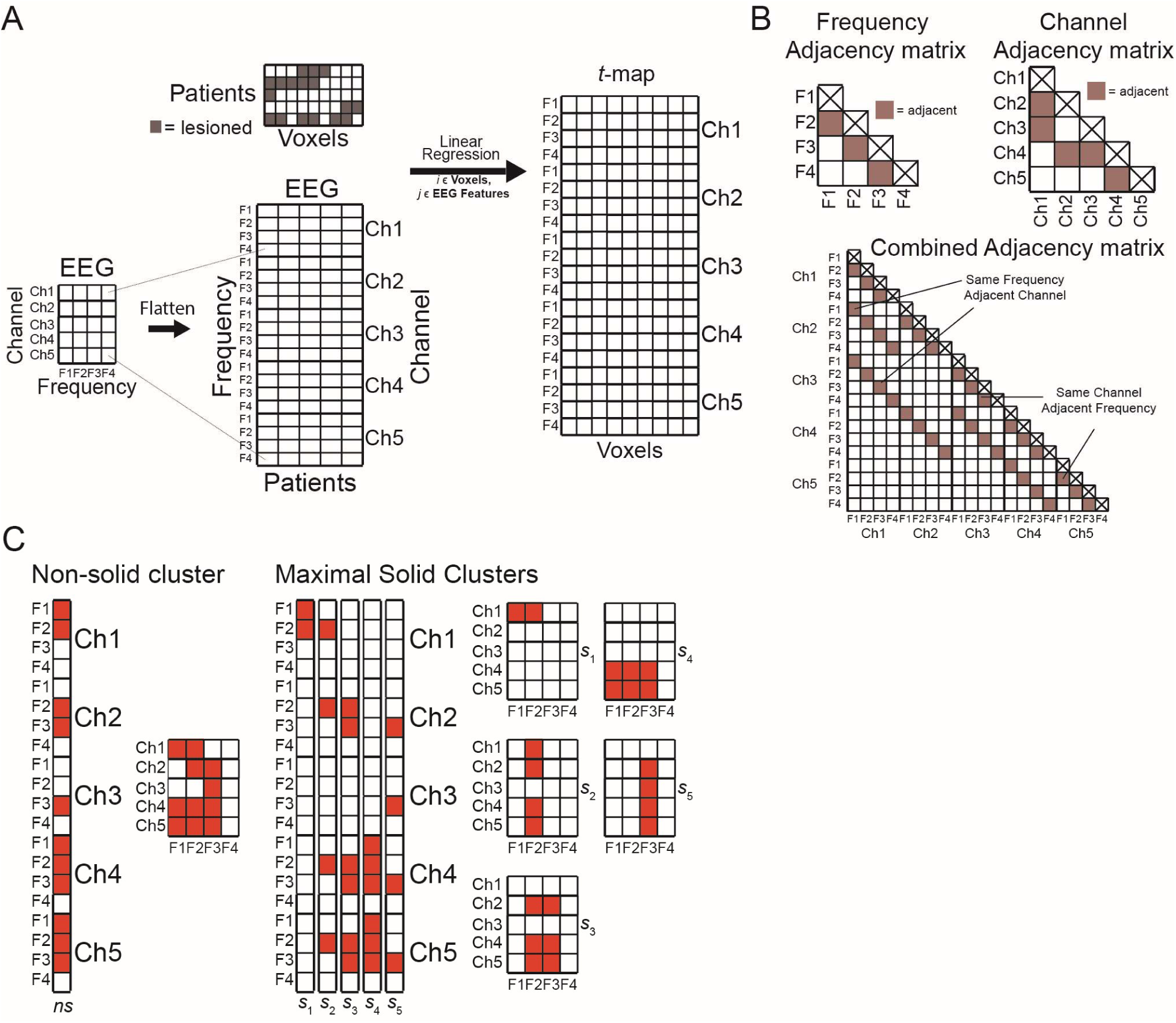
Additional steps to relate N-dimension EEG data to lesions. **(A)** In the N-dimensional EEG data case, each patient will have a set of N-dimensional EEG features (e.g., EEG [channels, Frequencies]) (Fig. 2A). Statistics are again applied for each combination of EEG Feature and voxel. **(B)** An adjacency matrix needs to be defined for each dimension (in this case, Channels and Frequency). A combined adjacency matrix can be created from these, where the same frequency in adjacent channels, and the same channel in adjacent frequencies will be defined as adjacent. **(C)** Resulting maximal EEG feature combinations are a combination of N-dimensional tuples. These can be non-solid, meaning that not all combinations features along each dimension are present. This can make interpretation of clusters complex. Non-solid clusters can be decomposed into maximal solid clusters (Fig. S1), where combinations of features are present for all dimensions.

**Figure 5).**
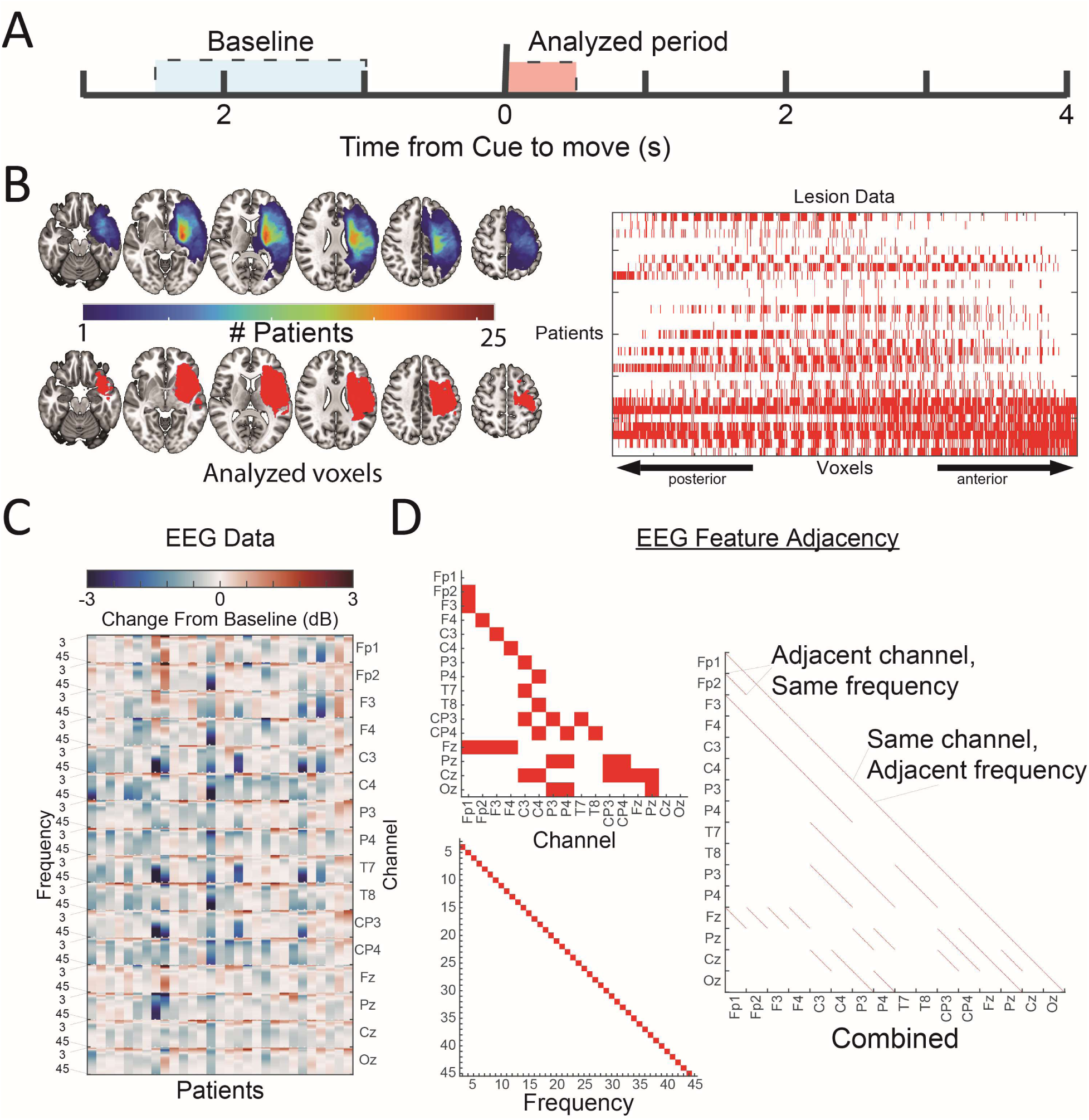
Input data for EEG-VLM from cued movement task in chronic stroke patients. **(A)** 29 chronic stroke patients (ischemic MCA) with hand paresis performed attempted cued hand movement for 4 seconds. (Reanalysis of data from (López-Larraz et al. 2018)**) (B)** (left) lesion overlay maps for 29 subjects, and voxels used for lesion mapping which have >= 5 subjects with lesions. (right) flattened lesion maps for all 29 patients for the analyzed voxels. **(C)** Baseline corrected power changes (Mean of Analyzed period for 16 channel EEG at 3 to 45 Hz at 1 Hz intervals. **(D)** Adjacency maps for EEG features.

**Figure 6).**
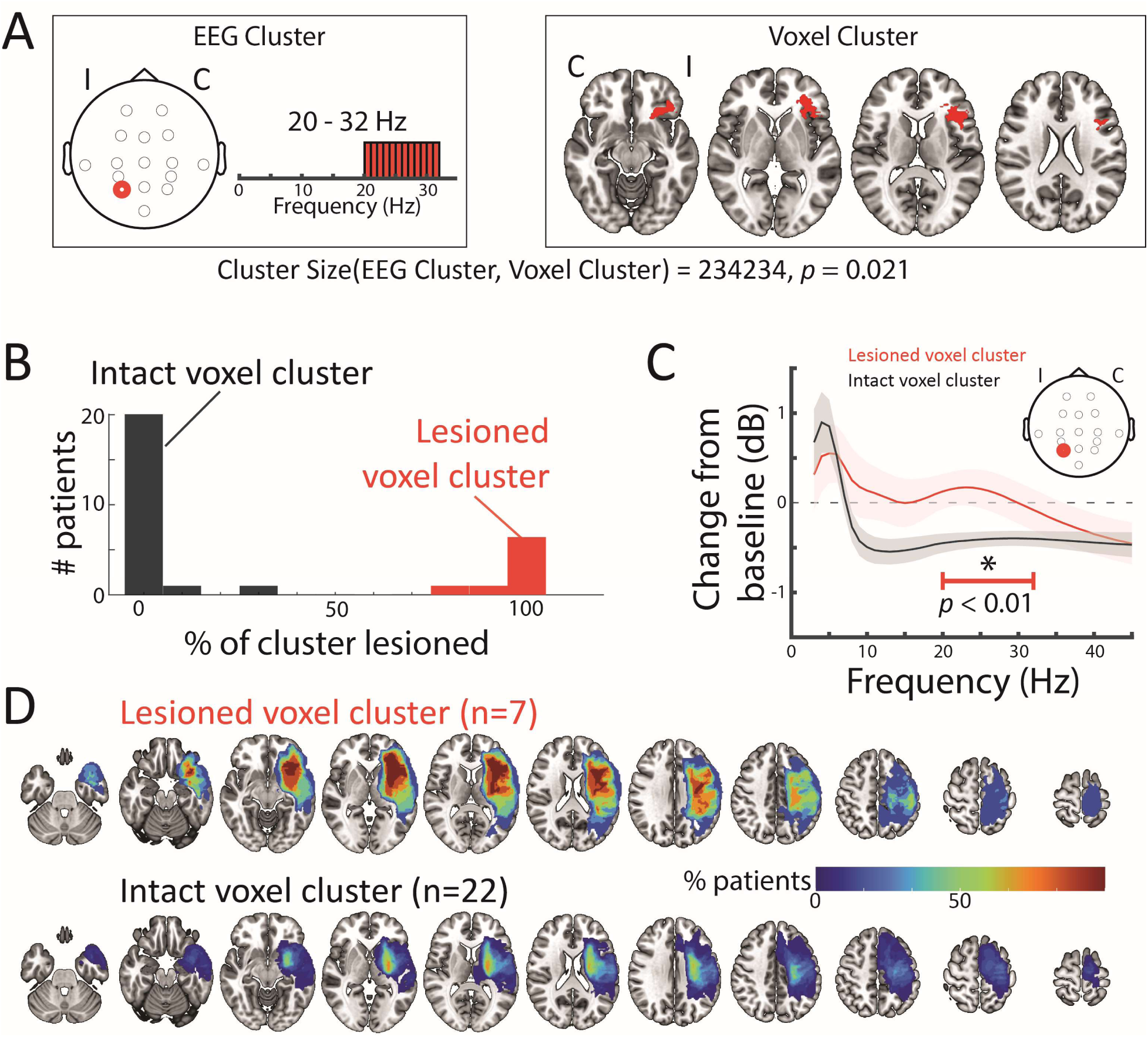
Frontal white matter lesions correlate with reduced ipsi-lesional parietal cue-evoked responses. **(A)** EEG-VLM found a significant EEG feature-lesion cluster consisting of frequencies 20-32 Hz over the ipsilesional parietal (P3) electrode, related to a frontal white matter lesion cluster. **(B)** Histogram of percent of the lesion cluster from (A) that is lesioned for each patient. For illustration purposes in (C) and (D) patients were split into 2 groups based on whether more than 50% of the voxel cluster was lesioned. **(C)** EEG changes for the lesioned vs. non-lesioned group electrode P3 show greater beta (20-32 Hz) desynchronization in the non-lesioned group. **(D)** Lesion overlays for the 2 group show that the lesioned group lesion extends more anterior. (I = Ipsilesional, C=Contralesional)

## Methods

We started from an implementation of voxel-lesion symptom mapping using cluster permutation statistics (Maris and Oostenveld 2007), where the input is zero-dimension (scalar; 0D, corresponding to a single value per patient) EEG data (Fig. 2). We then extended this method to 1-dimensional (1D, corresponding to several EEG values per patient e.g. one value for every channel) data (Fig. 3) and, further, to N-dimensional data (ND; Fig. 4). The overall aim is to find the correlations between clusters of EEG features and lesioned voxel-clusters that are largest (i.e., where size is determined by summing significant test statistics across all EEG-feature voxel pairs), such that a cluster with the same amount of voxels but more EEG features is larger than one with less EEG features, a cluster with the same amount of features and more voxels is larger than one with less voxels, and a cluster with the same amount of features and voxels and more significant correlations is larger than one with less significant correlations. As a smaller number of EEG features can be associated with a larger lesioned voxel cluster (Fig. 1D) and vice versa, the method needs to be capable of finding all possible relationships. However, looking at all possible combinations of EEG features would be intractable for a large numbers of features (e.g., with 100 EEG features that are all adjacent there would be ∼10^30^ possible EEG clusters). Thus, we developed an algorithm that detects the largest clusters present in the data without needing to check every combination.

### Zero-dimensional (0D, Scalar) Voxel-based Lesion Symptom Mapping (VLSM) 0D: Input data

For all forms of VLSM, lesion maps are required for each patient. These consist of a set of voxels (in a common coordinate space (e.g., Talaraich, MNI, etc. (Evans et al. 2012)) which covers the entire brain and indicates whether or not there is a lesion present at that voxel. For statistics to be performed at a voxel, there needs to be a large enough set of subjects that do and do not have a lesion at that voxel (e.g., N>=5 do have a lesion and N>=5 do not have a lesion (Karnath, Sperber, and Rorden 2019)). The other input data required is EEG data. For the 0D case, EEG data need to be reduced to a single value for each subject, for example by averaging beta power over motor electrodes.

### 0D: Statistics

The relationship between lesion presence and the EEG value is assessed at every voxel, to give a statistic at every voxel. Standard practice is to use linear regression (see methods, Equation 1), with *t*-statistics extracted (Equation 2). If required, confounders (such as overall lesion size, age, sex) can also be added (Equation 3).

### 0D: Clustering

Due to the large number of statistical comparisons (equal to the number of voxels), statistics need to be corrected for multiple comparisons. This can either be done at the voxel level (e.g., using Bonferroni correction or False Discovery Rate (FDR)), or clustering can be applied, and statistics done for each cluster. Here we apply a clustering procedure (Maris and Oostenveld 2007), based on adjacency of voxels (Fig. 2B). In the first step, the t-map is thresholded at a set significance level (the cluster defining threshold, alpha (e.g., *p*<0.05)). Significant voxels (with same sign t-statistic) that are adjacent to each other are then formed into clusters. Each cluster is assigned a cluster size, which is a summary statistic equal to the sum of the t-statistics from all voxels within the cluster. By assigning a size to each cluster, we can then compare the relative size of different voxel clusters.

### 0D: Permutation

To calculate the significance of each cluster, we compare the cluster size to a null distribution, which consists of the maximum cluster sizes that you expect to see by chance. This distribution is calculated by permuting the data multiple times (e.g., 1,000 times). For each permutation, the link between EEG and lesion maps is broken by shuffling the subject labels of the EEG data (Fig. 2C). After shuffling, statistics (Fig. 2A) and clustering (Fig. 2B) are applied in the same manner as was done for the unshuffled data. The maximum cluster size is then taken for each permutation to form the null distribution (Fig. 2D). Taking the maximum cluster size across all permutations forms the null distribution, and the original cluster sizes are compared to this. Clusters in original data with summary statistics exceeding the 95th percentile of null distribution are significant (corresponding to *p* < 0.05, cluster-corrected, one-tailed test).

### One-dimensional (1D) Voxel Lesion Symptom Mapping (VLSM)

For one-dimensional EEG data, there are additional steps that are required (Fig. 3A) compared to the 0D EEG data. These steps (Steps 2 and 3) are necessary to find the EEG clusters (which are defined sets of connected EEG features) that are present in the data (meaning that for at least one voxel, all EEG features in the cluster have a t-statistic > threshold) and maximal (meaning that for all voxels significant for this EEG feature cluster it is not possible to add another additional EEG feature to the cluster that would be significant for all of the voxels).

### 1D: Input data

For 1D data (Fig. 3B), the input data consist of the same lesion map data as required for 0D (scalar) analysis, but the EEG data now have *X* values per patient. Depending on the analysis, these *X* values could represent different EEG frequencies, timepoints or channels (as shown in Fig. 3B). EEG data naturally have spatio-temporal correlations (e.g., autocorrelations in time, volume conduction between nearby channels, correlations between adjacent frequencies). Therefore, it is usually not possible to consider each of the X values as independent. Cluster based permutation analysis (Maris and Oostenveld 2007) addresses this, by forming clusters of significance in EEG values data. Using adjacency matrices, which assign the values that are reasonably close to be considered as a single item. For example, in the spatial dimension, nearby channels would be considered adjacent (Fig. 3C). Therefore, for 1D-VLSM, an adjacency matrix, is an additional component required for the EEG dimension, as compared to 0D-VLSM.

### 1D: Statistics

In Step 1 (Fig. 3B), the relationship between lesion presence and EEG feature is assessed for every [voxel, EEG feature] pair resulting in a *t*-map with dimensions (#voxels x #EEG features) using the same statistics as were used for the 0D case. (methods, Equation 1,2,3).

### 1D: Clustering

Compared to the 0D case, we now have a *t*-map for every EEG feature. Due to spatial and temporal dependencies inherent in the data, these features (t-maps) cannot be treated as independent. One method to address this is to combine significant results across EEG features that are adjacent to form clusters, and then find the voxels where all the EEG features in the cluster are significant. This will result in EEG clusters (set of adjacent EEG features, e.g., two adjacent occipital EEG channels) that are significant for a specific lesion cluster (set of adjacent voxels). Therefore, to make inferences about the resulting t-map, it is easier to split clustering into a three-step process: first define the adjacency between EEG features (Fig. 3C), then find the maximal EEG clusters (Fig. 3D), and finally cluster the voxels for which every feature in the EEG cluster is significant (Fig. 3F).

### Finding maximal EEG clusters

With *n* EEG features there are 2^n^ – 1 possible combinations of these features (Fig. 3C). However, some of these combinations will not form a connected cluster (meaning that all features in the cluster can be connected through the adjacency matrix, e.g., channels 1, 2 and 4 would form a connected cluster as 1 is connected to 2 and 2 is connected to 4). With large numbers of EEG features it quickly becomes computationally intractable to assess each combination (e.g., with a 128 channel EEG net, there would be 2^128^-1 possible combinations of channels). The number of possible connected clusters depends on the EEG feature adjacency matrix, and the set of possible connected clusters will be smaller than the set of all combinations (if all features are not adjacent), but often will nonetheless be computationally intractable with large numbers of features. The set of EEG clusters that we are interested in (those that are present in the thresholded t-map and maximal) will be much smaller than the set of connected clusters, therefore identifying this set in an efficient manner can dramatically reduce the computational load and make the problem tractable.

Therefore, in Step 2ii, the aim is to find all **maximal** EEG clusters that are **present** in the thresholded t-map. By maximal we mean that this EEG cluster is not a subset of another EEG feature cluster, which is significant for all the same voxels. While there are usually thousands of voxels present in the t-map, there will be many voxels that have the same EEG features that are significant. As such, we can reduce the size of the problem by considering unique combinations of EEG features (Fig. 3Dii). While each of the feature combinations may be present, it is possible that the features are not adjacent (e.g. although there are voxels significant for channels 1, 4 and 5 (Fig. 3Di,ii); these do not form an EEG cluster as channel 1 is not adjacent to channel 4 or 5). Therefore, in the next step (Fig. 3Diii), each of the unique feature combinations is clustered using the adjacency matrix (Fig. 3C). As some of these clusters will be common across the combinations, the set of unique maximal EEG clusters is formed (Fig. 3Div). Note that while step 3Dii is not strictly necessary, it does reduce the computational load of the problem, as each combination of EEG features only needs to be clustered once.

### Clustering voxels for each EEG feature cluster

We can now find which voxels are associated with each EEG feature cluster (Fig. 3E) by identifying which voxels are significant for every feature in the EEG feature cluster. We can then cluster these voxels in 3D space (as in Fig. 2B) and calculate a cluster size. The cluster size is summed across all pairs of EEG features in the EEG feature cluster and voxels in the lesion cluster (Methods, Equation 4).

### N-Dimensional (ND) Voxel Lesion Symptom Mapping (VLSM)

### ND: Input data

For *N*-dimensional data (Fig. 4A), the input data consist of the same lesion map data as required for 0D and 1D analysis, but the EEG data now have *X x Y x…* features per patient, where each X, Y,… represents a different dimension of EEG data. For example, X could represent different channels and Y could represent different frequencies (as shown in Fig. 4A). Each dimension will have its own associated adjacency matrix. All EEG features can be collapsed into a single dimension and adjacency matrices combined.

### ND: Statistics

As in the 1D case, the relationship between lesion presence and EEG is assessed for every [voxel, EEG feature] pair resulting in a t-map with dimensions (#voxels x #EEG features).

### ND: Clustering

As in the 1D case, this is a two-step process with the second step (clustering voxels) being identical to the 1D case. For the first step, it is required to define an adjacency matrix for each EEG dimension (in this case, for frequency and channel). As these ND data can be flattened to a single dimension for each patient (as seen in Fig, 4A), it is possible to combine the adjacency matrices for each dimension (Fig. 1B). This puts the data into a form where the same method as 1D data can be applied.

However, when finding maximal EEG feature clusters, it may be preferable to add an additional step that refines the EEG feature clusters. When there is more than 1D, it is possible after finding the unique maximal EEG feature clusters that the EEG feature cluster contains a scattered mix of feature combinations across the different dimensions. Here we introduce a property of an EEG feature cluster, which is whether it is “solid”. In a non-solid EEG feature cluster, although there is adjacency between all EEG features, it is not the case that all combinations across the N (X, Y, …) dimensions are present. For a solid EEG feature cluster, all combinations will be present. For example, in Figure 4B, frequencies F1, F2 and F3 are significant in the non-solid cluster for C5, but only F3 is significant for channel C3. This limits the interpretability of the EEG feature cluster, as rather than just knowing the features along each EEG dimension, you will need to know all pairs that are present, changing the amount of info to remember from X+ Y+ … to X*Y*… Therefore, we introduce a method that decomposes non-solid clusters into maximal solid clusters, which can be defined by a set of values for each dimension, where all values in that dimension are adjacent and all tuples across the dimensions are significant. Figure S1 describes the additional steps necessary to decompose a non-solid cluster into its constituent maximal solid clusters.

### EEG-VLM algorithm

The EEG VLM algorithm was developed using custom-written code in MATLAB.

### Required Input Data

Required data for the EEG-VLM algorithm are (1) lesion maps for each patient in a common MNI standard stereotaxic space, (2) N-dimensional EEG data for each subject and (3) when the number of dimensions of the EEG data is greater than 0, then an adjacency matrix is required for each dimension (Fig. 3B).

### Dimensions in EEG data

The algorithm can be applied to EEG data that has >= 0 dimensions. Zero-dimensional (0D) EEG data corresponds to the case where there is a single EEG feature per subject. For example, this single value could be calculated by averaging the power in a frequency-band across a selected set of channels. One-dimensional (1D) EEG data corresponds to the case where there are multiple EEG features per subject for a single dimension. For example, if the dimension is EEG channels, then each subject would have a value for each of the EEG channels. Two-dimensional (2D) EEG data would mean that each subject would have a value for each combination of the 2 dimensions. For example, if the dimensions were channels and frequency, then the subject would have a value for each combination of EEG channel and EEG frequency. Additional dimensions such as time could also be added for N-dimensional (ND) EEG data.

### Properties of EEG feature clusters

#### Connected cluster

A connected EEG feature cluster indicates that all features in the cluster can be connected through the EEG feature adjacency matrix, e.g., in Figure 3C channels 1, 2 and 4 would form a connected cluster as channel 1 is connected to channel 2 and channel 2 is connected to channel 4.

#### Maximal cluster

Here we define a maximal EEG feature cluster (that is significant for a set of voxels), as a cluster that is not a subset of another EEG feature cluster, which is significant for all the same voxels.

#### Solid clusters

Here we define a property of EEG feature clusters, which is whether they are solid. To determine if an EEG feature cluster is solid, we follow this algorithm:

1. For each EEG dimension, determine the set of EEG features present
2. A solid cluster will contain all combinations across the sets of present EEG features for each dimension.

For example, for 2D EEG data, if the set of EEG features present in a cluster for the channel dimension was {Oz, Pz} and the set of frequencies was {19, 20} Hz, then the solid EEG feature cluster would consist of [Oz, 19], [Oz, 20], [Pz, 19], [Pz, 20]. An EEG feature cluster consisting of [Oz, 20], [Pz, 19], [Pz, 20] would be non-solid as it is missing one of the possible combinations [Oz, 19]. Note that all EEG features clusters with zero and one dimension are solid.

### Steps of algorithm

The EEG VLM algorithm follows four steps (Fig. 3C):

### Step 1: Statistics

Linear regression is applied for every pair of [voxel_i_, EEG Feature_j_] (Equation 1), and the resulting *t*-statistic for the EEG predictor can be calculated according to (Equations 2 and 3).

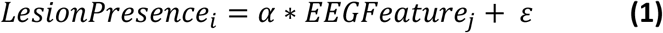

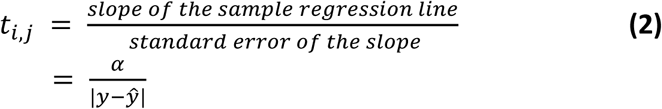

Confounds such as lesion size can also be included in the linear regression, e.g.,

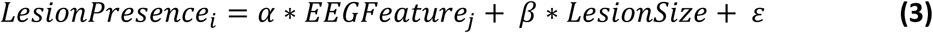

### Step 2: Clustering

Clustering is a 3-part process where **step 2i** defines how EEG features cluster **step 2ii** finds the maximal EEG feature clusters present in the data and **step 2iii** extracts the significant voxel clusters for each map. Steps 2ii and 2iii use a thresholded t-map, which is defined according to the *t* statistic exceeding a critical t value, which is calculated according to required p-value and degrees of freedom. The EEG VLM algorithm aims to aggregate all relationships within the thresholded t-map into a set of tuples [maximal solid EEG feature cluster, voxel cluster]. Note that a single significant relationship within the t-map (i.e., [single EEG Feature, single voxel]) can appear in multiple of these tuples. For example, this single relationship may appear in one tuple that contains a small number of EEG features in the EEG feature cluster, and a large number of voxels in the voxel cluster, while the single relationship also appears in another tuple that has a large number of EEG features in the EEG feature cluster but only a small number of voxels in the voxel cluster (Fig. 1C).

### Step 2i: Define EEG feature adjacency

In the case where the EEG data has more than zero-dimensions (0D) (i.e. more than one EEG feature per patient), it is necessary to define an adjacency matrix for each EEG dimension (Fig. 3C, 4B). These adjacency matrices are usually defined based on spatiotemporal correlations inherent in the data. For example, due to volume conduction, nearby EEG channels usually show similar activity, which can be addressed using a measure of adjacency defined using Euclidean distance between the channels. For spectral or temporal features, adjacency can be defined by temporal or spectral distance (e.g., time bins next to each other are adjacent, and frequencies next to each other are adjacent).

### Step 2ii: Find EEG clusters

This details of performing this step depend on the number of dimensions in the EEG data. For zero-dimensional (0D) data, there is only one EEG feature in the t-map and so only one possible cluster. For one dimensional (1D) data, the resulting t-map can be clustered according to that EEG dimensions adjacency matrix (Fig 3C), according to the following algorithm:

1. Threshold the t-map according to the critical t-value
2. Find all unique combinations of EEG features that are present in the thresholded t-map.
3. For each combination, cluster the EEG features. This produces a set of EEG feature clusters for each combination.
4. Combine all sets of EEG feature clusters and remove any that are duplicates.

For N-Dimensional data, the clustering performed in step 3 uses a combined EEG adjacency matrix. An additional step is then performed which transforms any of the found EEG feature clusters that are non-solid, into a set of maximal solid feature clusters according to the follow algorithm (Fig. S1):

1. Input: non-solid *N*-dimensional EEG feature cluster
2. For each dimension (X):

a. Reshape the input cluster into a 2D matrix [dimension (X), all other dimensions]
b. Find all unique combinations of dimension(X) in this matrix and cluster each combination using the adjacency matrix for dimension X. (Fig. 4Ci)
c. For each found cluster (Y), check for hidden clusters, which are smaller clusters that span multiple combinations (Fig. 4Cii):

i. Mask each combination by cluster Y (i.e., only keeping the features for that combination that are in Y).
ii. Cluster the combinations again and see if there are any hidden clusters that were not previously found.
d. Repeat step 2c until no new clusters are added, giving a set of clusters for dimension X (**C_x_**).
e. Compare all clusters in **Cx** and determine which ones are subsets of each other (Fig. 5Ciii).
3. Check all possible N-tuples of clusters (i.e., one from each dimension) to see which ones are present in the input.
4. For each N-tuple that is present:

a. Check if it is a subset in each dimension of another present N-tuple; if it is remove it.
5. The remaining N-tuples are maximal solid EEG Feature clusters (Fig. 4Cv).

### Step 2iii: Find Voxel clusters

For each EEG feature cluster, determine the voxels that are significant for every member of the EEG Feature Cluster. These voxels can then be clustered in 3D space according to the 3D space adjacency matrix. (Fig. 5E).

The cluster size for each pair [EEG feature cluster, voxel cluster] can then be calculated according to equation 4:

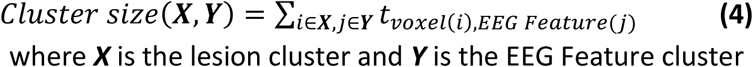

### Step 3: Permutation

To calculate a null distribution of cluster sizes, a large number of permutations (e.g., 1,000) are run, where for permutation the subject label for the EEG data is permuted (Fig. 2C). Steps 1 and 2 are then run for each permutation, and the largest cluster sizes showing a positive and a negative correlation are taken. This forms a null distribution.

### Step 4: Significance

Cluster sizes from the unshuffled analysis are compared to the null distribution and are significant if they are larger than 95% of the null distribution (equivalent to a p-value of < 0.05) (Fig. 3D).

### Task

The EEG task and dataset was previously described in (López-Larraz et al. 2018), with relevant details reproduced here.

### Patients

Thirty-one chronic stroke patients (19 male, mean age 54.0 ± 11.7 years, range 29–73, time since stroke 60.3 ± 58.2 months, range 10–232) were recruited for this study. Inclusion criteria were: (1) hand paralysis with no finger extension; (2) minimum time since stroke 10 months; (3) age between 18 and 80 years; (4) no psychiatric or neurological condition other than stroke; (5) no cerebellar lesion or bilateral motor deficit; (6) no pregnancy; (7) no epilepsy or medication for epilepsy during the last 6 months; (8) eligibility to undergo magnetic resonance imaging (MRI); and (9) ability to understand and follow instructions. Demographic and clinical data of the patients can be found in Table 1. Further details about the lesion location and the brain areas affected by the stroke in each patient can be found in Supplementary Table 1 and Supplementary Fig. 1. The experiments were conducted at the University of Tübingen, Germany. The experimental procedure was approved by the ethics committee of the Faculty of Medicine of the University of Tübingen, and all the patients provided written informed consent.

### Experimental design and procedure

The patients were asked to perform an assessment task in which they had to move their healthy hand, or to attempt to move their paralyzed hand, while their EEG and electromyographic (EMG) activity was recorded. Each patient performed between 4 and 6 blocks, each of which contained, in a random order, 17 trials of movement execution (healthy hand) and 17 trials of movement attempt (completely paralyzed hand). The target number of blocks for patients was 6, though the actual number of blocks recorded depended on patient fatigue levels, as in some cases they requested to stop the measurements before finishing the 6 expected blocks. Audiovisual cues guided the patients regarding the phases of a trial: rest (random duration between 4 and 5 s), movement execution/ attempt (4 s), and an inter-trial interval that was included every three trials (random duration between 8 and 9 s). The patients were asked to perform (or try to perform) openings and closings with their healthy (or paretic) hand at a comfortable personal pace during the 4 s of the movement execution/attempt interval (generally, it took them around 1.5 s to perform a complete open-close cycle). Before starting the first block, they were instructed to minimize compensatory movements and any other source of contamination for the EEG activity during the rest and movement intervals.

### Data recording and preprocessing

EEG activity was recorded using a commercial Acticap system (BrainProducts GmbH, Germany), with 16 electrodes placed on Fp1, Fp2, F3, Fz, F4, T7, C3, Cz, C4, T8, CP3, CP4, P3, Pz, P4, and Oz (according to the international 10/20 system). The ground and reference electrodes were placed on AFz and FCz, respectively. Vertical and horizontal electrooculography (EOG) was also recorded to capture eye movements. Electromyographic activity was recorded using bipolar Ag/ AgCl electrodes (Myotronics-Noromed, USA) from four muscle groups of each arm: extensor carpi ulnaris, extensor digitorum, external head of the biceps and external head of the triceps. All signals were synchronously recorded at 500 Hz. Ocular artifacts were removed, (as previously described in (López-Larraz et al. 2018). The EEG data were filtered between 0.1 and 48 Hz with a 4-th order causal Butterworth filter. The signals were trimmed down to 7-s trials, from −3 to +4 s with respect to the movement cue. Although the shortest duration of the rest intervals was 4 s, we decided to consider only the last 3 s to ensure that the patients were in resting state. Data was then re-referenced using a Laplacian montage using the nearest neighbors for each electrode to reduce volume conduction recorded by neighboring electrodes. Trials were statistically rejected based on EMG and EEG as previously described in (López-Larraz et al. 2018)

### Time-frequency analysis

Time-frequency analysis was applied using the fieldtrip toolbox (Oostenveld et al. 2010), to extract trial-averaged power in frequencies from 3 to 45 Hz in 1 Hz bins, at a time resolution of 0.01 seconds using Morlet wavelets (3 cycles). Trial averaged power was then baseline corrected using the decibel method and the baseline period -2.5 to -1 second from the cue onset. To look at cue-evoked neural responses, the time-period from 0-0.5 seconds from the cue was averaged, giving EEG data for each subject with dimensions 16 channel x 42 frequencies.

### EEG Feature Adjacency Matrices

EEG Channel adjacency matrix (Fig. 5B) was defined based using Euclidean distance between electrodes. The EEG frequency adjacency matrix was defined by consecutive frequencies being adjacent.

### Lesion Maps

The method for extraction of the lesion masks was previously described in (Caria et al. 2020) and is summarized here. MRI recordings were performed with a 3-T Magnetom (Tim Trio, Siemens, Erlangen) whole-body scanner. A T1-weighted anatomical MR image was acquired using a 1-mm isotropic MPRAGE sequence with the following parameters: TR = 2300 ms, TE = 3.03 ms, TI = 1100 ms, flip angle = 8°, FOV = 256 × 256, matrix size = 256 × 256, number of slices = 176, slice thickness = 1 mm, bandwidth = 130 Hz/Px. Individual lesion masks were created manually on axial slices of a T1-weighted images using the MRIcron software package (see supplementary figure 2). Normalization to standard space was accomplished using the SPM8 normalization function.

### EEG_VLM settings for analysis

EEG VLM was applied using lesion maps, cue-evoked EEG activity, and adjacency matrices. Statistics for each pair [voxel, EEG Feature] included a lesion size confound. Each t-map was thresholded using a cluster-defining threshold equivalent to *p*<0.005. A total of 1,000 permutations were run to form the null distribution of cluster sizes and positive or negative cluster sizes were defined as significant if they were larger than the 95^th^ percentile of the corresponding null distribution.

## Results

### Application of method to EEG recordings in chronic stroke patients

Twenty-nine chronic stroke patients attempted movement with their paretic hand while their neural activity was recorded using a 16-channel, Acticap EEG system (see *Methods*). Each trial consisted of a 4 second inter-trial period followed by an audiovisual cue (Fig. 5A), after which subjects attempted to move their paretic hand. In control trials (which were analyzed separately) subjects were instructed not to move in response to the cue. Lesion maps were extracted from MRI scans, mapped into a common MNI space, and patients with lesions on the right hemisphere were flipped onto the left (Fig. 5B). EEG was recorded and analyzed from 16 channels, and to allow group comparison of neural activity on the ipsi- and contra-lesional hemisphere, patients with right-hemisphere lesions had left/right EEG channels flipped. After standard EEG preprocessing (see *Methods*), average wavelet power in frequencies from 3-45 Hz in the half second after the cue was calculated and was then baseline corrected (dB method) using the period -2.5:-1 second from the cue (Fig. 5C). These time periods were selected as they allow comparison of cue-evoked activity in the attempted movement trials as well as the control (no attempted movement) trials and will be uncontaminated by movement-related EMG activity. Adjacency maps for EEG channels (based on distance) and frequencies were defined (Fig. 5D).

The EEG-VLM method was applied to this dataset, using the lesion maps, 2D EEG data and adjacency matrices. For the statistics, lesion size was included as a confounder. This resulted in 3,775 EEG-VLM clusters showing a positive relationship between lesion presence and EEG (i.e., subjects with a lesion overlapping with the lesion cluster tended to have higher values for those features than subjects who didn’t), and 3,425 EEG-VLM clusters showed a negative relationship.

Testing significance of the cluster sizes showed that eleven of the clusters showing a positive relationship were significant but none of the EEG-VLM clusters showing a negative relationship were. Of the 11 positive clusters, all were related to a single channel (P3), and the frequencies were largely overlapping, with the largest range being 20-32 Hz and all the clusters including 23-28 Hz. Voxels for these 11 clusters were also highly overlapping, sharing at least 65% of the same voxels. The EEG-VLM with the largest cluster size (Fig 6A), consisted of frontal white matter lesion (1,291 voxels), with lesions in this cluster positively related to 20-32 Hz EEG evoked activity in electrode P3, which overlies the ipsilesional parietal cortex. Investigating the subjects overlap with this lesion (Fig. 6B), there was a clear bimodal distribution with most subjects (22) overlapping < 50 % of the lesion, and 7 subjects showing almost 100 % overlap with the lesion. Comparing the EEG data between the two groups showed that subjects without a lesion exhibited a broad beta desynchronization in this post-cue period, whereas subjects with the lesion showed little change from baseline during cued movement, suggesting an impaired cue-evoked response in subjects with this frontal-white matter area lesioned. Finally, the lesion overlap maps showed that the lesioned group tended to have infarcts that spread more anterior than those in the non-lesioned group. To evaluate whether this effect was related to the cue or to the movement, the same cluster was tested in the control condition, where subjects did not have to move after the cue. Here, the cluster showed a significant effect in the same direction (p=0.003), suggesting that this effect was in response to the cue rather than a movement-related finding. Taken together, these results suggest impaired cue-evoked responses in frontal-parietal networks when this frontal white matter region is damaged.

## Discussion

Here we introduce a method (EEG-VLM) for extending voxel-lesion mapping to multi-dimensional EEG data, which clusters all relationships between lesion presence (in voxels) and EEG features and defines the significance of those clusters. As proof of principle, we applied this method to lesion-map data from a group of chronic stroke patients who had EEG recorded while attempting hand movements with their paretic hand (Fig. 5). We found significant relationships between lesion presence and first 500ms after imperative cue evoked EEG activity (Fig. 6), with lesion clusters located in frontal white matter regions related to less beta desynchronization (20-32 Hz) in an EEG electrode overlying lesioned parietal cortex. This relationship was also significant in a control condition where subjects were instructed not to move in response to the cue, suggesting that this relationship was not specific to movement.

Despite decades of EEG research in patients with stroke, there has not been a concerted effort to understand how lesion location affects neural activity in these patients. In animal models, the effects of induced lesions on different areas on EEG activity has long been studied (Ball, Gloor, and Schaul 1977), and have been used to determine the genesis of cortical activity such as specific neural oscillations. However, it remains unclear how these findings translate to stroke-related infarcts in humans, among whom lesion patterns differ substantially from those induced experimentally in preclinical models (Taha et al. 2022). One classic study (Macdonell et al. 1988) investigated the potential of EEG to dissociate cortical from lacunar infarction in humans during the acute phase post-stroke (< 2 weeks from infarction), and found that cortical strokes in the middle cerebral artery territory were more likely to produce lateralized abnormalities (as assessed qualitatively) in the EEG as compared to lacunar strokes, which rarely did. However, despite these early findings showing that lesion location differentially affects neural activity, there has been little follow-up. Other recent studies have also looked at the relationship between EEG and relatively coarse measurements of lesion location (Ray et al. 2017; Cassidy et al. 2020) or infarct volume (Wu et al. 2016), that do not include the fine-grained information in lesion maps. We propose that the ability to define accurate statistical relationships between lesion location and EEG abnormalities will enable an improved mechanistic understanding of the effect that stroke-related injury to different cortical and subcortical regions has on the genesis of post-stroke EEG signal changes. This knowledge could help understand EEG findings related to impairment and recovery seen in studies where structural imaging was not available (i.e., improved biomarkers (Boyd et al. 2017)), and provide potential therapeutic targets for neurorehabilitative intervention (e.g., closed-loop electrical stimulation) based on restoring deficits in a patient’s neural functioning as assessed by their EEG (Ramanathan et al. 2018).

While localization of function has a long and important history in neuroscience, there are other perspectives that need to be considered that emphasize that there is not always a one-to-one mapping between functions or features and a particular neural area (Noble et al. 2023). Neural degeneracy—the concept that multiple different neural areas can support a behavior—is particularly relevant, as it suggests that multiple areas receive the required inputs, and output onto the required downstream regions necessary to produce a required computation, even though this might not be the primary role of that area. This is particularly relevant in stroke, where it is proposed that recovery from impairment may occur through neural compensation (Makin and Krakauer 2023), where intact areas that have latent motor representations are upregulated in order to take over the required function or to engage other residual neural pathways. It remains unclear how this neural compensation would be reflected in EEG activity, e.g., if premotor areas assume the functional role from a damaged M1, it is unclear if neural oscillation responses from these premotor areas would start to resemble those of M1 as has been seen with BOLD activation in fMRI(Ward et al. 2007).

Here we have presented a new algorithm and toolbox that can relate multi-dimensional features (which can be clustered according to adjacency) to the presence of a lesion. There are several ways that this method can be extended to expand the set of questions that it can ask. The first pertains to use of lesions to calculate dysconnectivity relative to normal subjects. One criticism of VLSM is that the lesion maps used cannot account for similar (but not overlapping) lesions having similar effects, e.g., two different non-overlapping lesions on the corticospinal tract (CST) could cause the same areas to become disconnected, and lead to similar functional impairments. This has led to methods to estimate a “disconnectome” map (Thiebaut de Schotten, Foulon, and Nachev 2020; Griffis et al. 2021; Fox Michael D. 2018) from the lesion map, which estimates the cortical areas and white matter tracts that become disconnected due to the lesion, using normative databases of anatomical or functional connectivity in healthy subjects. These methods produce voxel-level maps, with each voxel representing the percent disconnection that each voxel has as a result of the lesion. This allows both remote and local effects of the lesion to be estimated. These maps can then be substituted for the original EEG maps in EEG-VLM without any modification of the algorithm, with the result highlighting the relationship of areas that become disconnected with the multi-dimensional EEG features.

A second way that EEG-VLM can be extended is by incorporating behavioral measures into the algorithm. For example, a motor impairment outcome measure such as upper-extremity Fugl-Meyer score (Gladstone, Danells, and Black 2002) could be included in the statistics (along with the EEG and lesion maps) using commonality analysis. This method has proved successful in relating models of relationships that examine how stimuli are represented in EEG and fMRI (Hebart et al. 2018; Flounders et al. 2019). In this method, the shared variance between each EEG feature, voxel and outcome measure is calculated. This requires subtracting the shared variance between the EEG features and lesion presence that is unrelated to the outcome measure. For example, if the outcome measure is a motor measure that is uncorrelated with cognitive outcomes, the result will be that the EEG-lesion relationships most correlated with the motor outcome measure will be identified, and not those related to cognitive outcomes. This requires adjusting the statistics in Step 1 (see methods) to calculate shared variance. As shared variance is a magnitude (i.e., only takes positive values), this will also not distinguish between positive and negative relationships, which would need to be assessed in a post hoc fashion. By incorporating behavioral measures, EEG-VLM will indicate which relationships between lesions and EEG are associated with a particular behavioral impairment. Finally, EEG-VLM currently requires the arbitrary specification of a cluster-defining threshold. Choosing more liberal thresholds (e.g., p<0.05) will result in clusters that are broader but contain less significant relationships, while stricter thresholds (e.g., p < 0.00001) will result in clusters that are smaller but contain stronger relationships. For cluster-based permutation methods, a threshold-free method has been developed (Smith and Nichols 2009) that gives a threshold-free cluster enhancement (TFCE) value for each voxel, which is calculated using all possible cluster sizes of which the particular voxel is part. Each voxel is then evaluated against the null distribution of maximal TFCE values obtained from permuted analysis. Adapting EEG-VLM to use threshold-free statistics requires additional steps. Firstly, the set of maximal EEG clusters needs to be calculated at each possible threshold and aggregated across all of them. A voxel-level map of TFCE values is then calculated separately for each cluster. To do this, instead of thresholding the t-statistic map for the present EEG features, each voxel instead takes the largest p-value (i.e., least significant) from the set of EEG features in the EEG feature cluster. This results in a vector equal to the number of voxels over which TFCE can be calculated in the usual manner.

When applying EEG-VLM, it is important to consider the sample size that is required to obtain robust results that are more likely to be reproduced in other samples (Marek et al. 2022; Spisak, Bingel, and Wager 2023). For VLSM methods (Lorca-Puls et al. 2018), it has been shown (through resampling from a larger distribution) that smaller sample sizes (*N*<=30) are generally underpowered leading to many relationships that are present but not being found, and for any significant results that are found, effect sizes will likely be overestimated. Larger sample sizes (*N* >=90) have been shown to have relatively stable effect sizes and are sufficiently powered for finding the present relationships. For multidimensional EEG data, power analysis for a certain cluster size will additionally depend on statistical aspects of the data, such as the correlation between adjacent features. Currently, EEG studies in stroke patients (especially if limited to acute time periods) are rarely that big, with most studies being <=30 subjects. The answer to this problem lies in data sharing and collaboration across institutions, with agreed upon experimental protocols and data standards (e.g. BIDS (Pernet et al. 2019)) being used to allow the required number of subjects to be aggregated. This data sharing practice has allowed large structural MRI datasets to be acquired from stroke patients, with the ENIGMA study recording over 2,100 patients across 39 research sites (Liew et al. 2022). While these caveats apply to the dataset used in this study, the evidence for the current results is boosted by the replication of the result in a separate control task. Rigorous experimental design can allow inferences to be drawn from smaller sample sizes.

We found post-cue changes in neural activity in patients with white matter PFC lesions over the lesioned parietal cortex. Studies have implicated lesions in this region with damage to the fronto-parietal attention network, leading to deficits in attention in these patients (Kaufmann et al. 2023). A recently published study (Raposo et al. 2023) reported that frontal lesions caused cue-related reductions in P100 amplitude over ipsilesional parietal channels (also seen in (Knight et al. 1981; Knight 1984; 1997; Voytek et al. 2010), which could be related(Li et al. 2018) to the evoked changes in beta-band oscillations found in the current study.

EEG-VLM is a novel and unbiased method for relating neurophysiologic changes after stroke with neuroanatomic lesions. We propose that this method will enable an improved mechanistic understanding of stroke-induced changes in EEG signals, improve detection of behaviorally relevant EEG-biomarkers, and guide closed-loop therapeutic stimulation approaches. Further, this method can be applied to other multi-dimensional datasets such as different neurophysiological recording methods (MEG, fNIRS) or kinematic recordings, to allow this data to be mapped to the presence of lesions without *a priori* selection of features of interest.

## Data and Code Availability Statement

Code and an example dataset were made available to reviewers when paper was submitted and will be available publicly on GitHub at the time of publication.

## Author Contributions

Richard Hardstone: Conceptualization, Methodology, Software, Formal analysis, Visualization, Writing – Original Draft, Writing – Review and Editing

Lauren Ostrowski: Investigation, Data Curation, Writing – Review and Editing

Aliceson N. Dusang: Investigation, Data Curation, Writing – Review and Editing

Eduardo López-Larraz: Resources, Investigation

Jessica Jesser: Resources, Investigation

Sydney S. Cash: Writing – Review and Editing

Steven C. Cramer: Writing – Review and Editing

Leigh R. Hochberg: Resources, Writing – Review and Editing

Ander Ramos-Murguialday: Resources, Investigation, Writing – Review and Editing

David J. Lin: Conceptualization, Writing— Review and Editing, Funding acquisition

## Acknowledgments

Analysis was performed on the MSLC Compute Cluster funded by the Massachusetts Life Sciences Center to support research at the Athinoula A. Martinos Center for Biomedical Imaging.

## Supplementary Material

**Figure S1).**
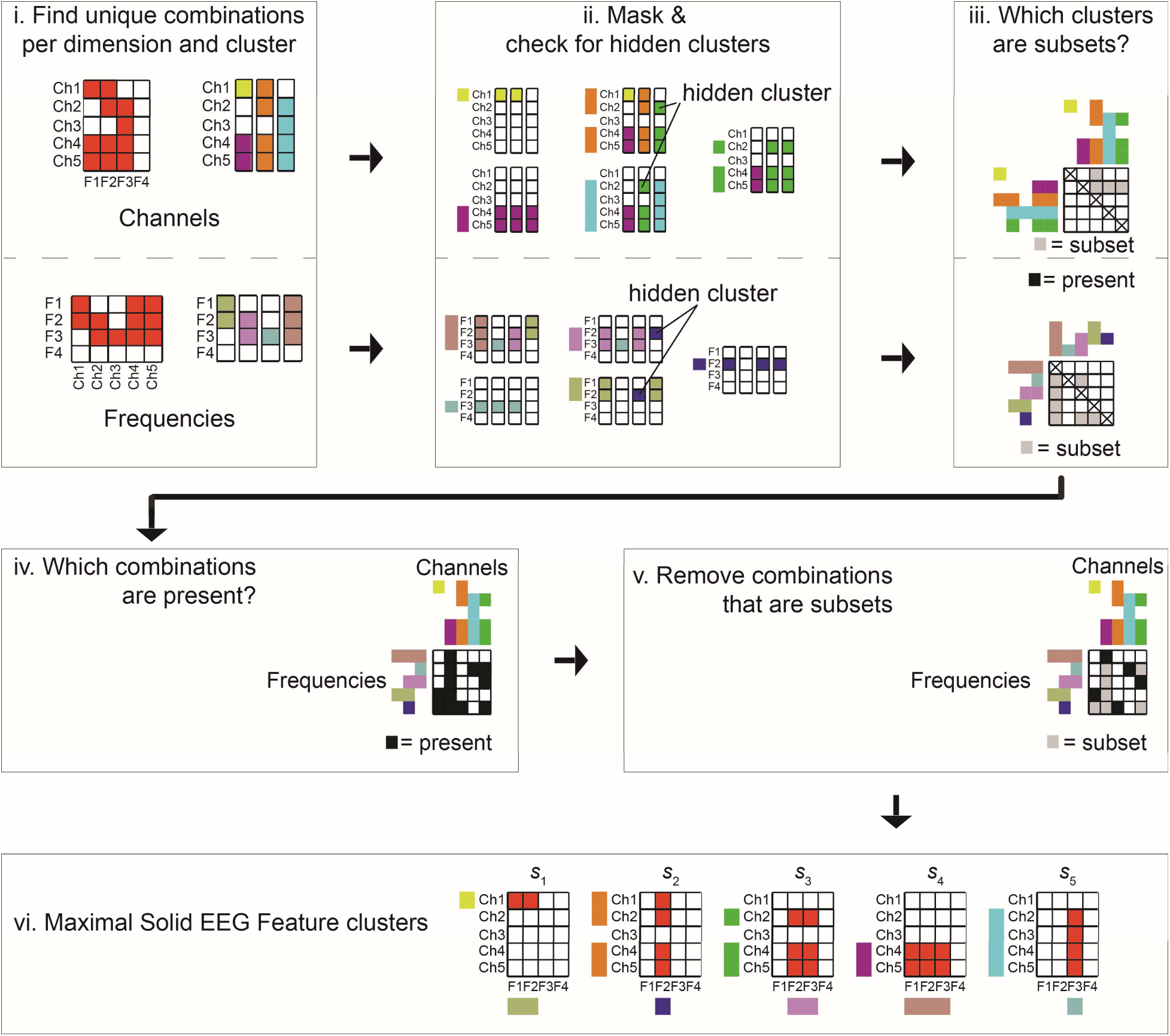
Frontal white matter lesions correlate with reduced ipsi-lesional parietal cue-evoked responses. Method to decompose non-solid *N*-dimensional cluster into maximal solid cluster. (i) For each dimension (In this case Channels and Frequencies), the unique combinations of features for that dimension are found and clustered. (ii) Each of these clusters is then checked to see if there are potential hidden clusters (clusters of features of that dimension that are present across multiple combinations but hidden by larger clusters). (iii) All clusters for this dimension are then compared to see which ones are subsets. (iv) All combinations of clusters across the N-dimensions are then checked to find which N-D combinations are present. (v) N-D combinations which are subsets for each dimension of another present N-D combination are removed (vi) This gives a set of maximal solid EEG Feature clusters which can then be clustered in 3D space (according to Fig. 3E).

### Supplementary Information

To make the problem tractable, each EEG feature dimension is first addressed individually to find all combinations that occur along that dimension. The cluster is reshaped into a 2D matrix consisting of (this dimension, all other dimensions). For each dimension we again find the unique combinations that occur, and then cluster each of those dimensions. We then need to check for “hidden” clusters which are masked by larger clusters in one of the other combinations. (*Note: hidden clusters will not definitely survive, as they are dependent on a cluster that spans multiple other features that may or may not be connected*). Therefore, after finding the set of present maximal clusters, for each member of this set, the present combinations are masked by that cluster. The masked data is then clustered again. This process continues, until no new clusters are found. The second step is to compare each of the clusters are work out which ones are a subset of each other. Once this has been done for each dimension, giving a set of clusters for each dimension, the algorithm then checks for the presence of all possible N-tuples of clusters across the N dimensions. After an exhaustive search of these has been completed, any clusters that are subset of a present cluster in all dimensions is removed. This gives the final set of maximal solid EEG Feature clusters.

## References

Ball, G. J, P Gloor, and N Schaul. 1977. “The Cortical Electromicrophysiology of Pathological Delta Waves in the Electroencephalogram of Cats.” Electroencephalography and Clinical Neurophysiology 43 (3): 346–61. 10.1016/0013-4694(77)90258-9.

Bates, Elizabeth, Stephen M Wilson, Ayse Pinar Saygin, Frederic Dick, Martin I Sereno, Robert T Knight, and Nina F Dronkers. 2003. “Voxel-Based Lesion–Symptom Mapping.” Nature Neuroscience 6 (5).

Benson, D. F., and D. H. Patten. 1967. “The Use of Radioactive Isotopes in the Localization of Aphasia-Producing Lesions.” Cortex 3 (2): 258–71. 10.1016/S0010-9452(67)80016-9.

Bentes, Carla, Ana Rita Peralta, Pedro Viana, Hugo Martins, Carlos Morgado, Carlos Casimiro, Ana Catarina Franco, et al. 2018. “Quantitative EEG and Functional Outcome Following Acute Ischemic Stroke.” Clinical Neurophysiology 129 (8): 1680–87. 10.1016/j.clinph.2018.05.021.

Bibián, Carlos, Nerea Irastorza-Landa, Monika Schönauer, Niels Birbaumer, Eduardo López-Larraz, and Ander Ramos-Murguialday. 2022. “On the Extraction of Purely Motor EEG Neural Correlates during an Upper Limb Visuomotor Task.” Cerebral Cortex 32 (19): 4243–54. 10.1093/cercor/bhab479.

Bönstrup, Marlene, Lutz Krawinkel, Robert Schulz, Bastian Cheng, Jan Feldheim, Götz Thomalla, Leonardo G. Cohen, and Christian Gerloff. 2019. “Low-Frequency Brain Oscillations Track Motor Recovery in Human Stroke.” Annals of Neurology 86 (6): 853–65. 10.1002/ana.25615.

Boyd, Lara A., Kathryn S. Hayward, Nick S. Ward, Cathy M. Stinear, Charlotte Rosso, Rebecca J. Fisher, Alexandre R. Carter, et al. 2017. “Biomarkers of Stroke Recovery: Consensus-Based Core Recommendations from the Stroke Recovery and Rehabilitation Roundtable.” International Journal of Stroke 12 (5): 480–93. 10.1177/1747493017714176.

Broca, Paul. 1861. “Remarques Sur Le Siège de La Faculté Du Langage Articulé, Suivies d’une Observation d’aphémie (Perte de La Parole).” Bulletin et Memoires de La Societe Anatomique de Paris 6:330–57.

Caria, Andrea, Josué Luiz Dalboni da Rocha, Giuseppe Gallitto, Niels Birbaumer, Ranganatha Sitaram, and Ander Ramos Murguialday. 2020. “Brain–Machine Interface Induced Morpho-Functional Remodeling of the Neural Motor System in Severe Chronic Stroke.” Neurotherapeutics 17 (2): 635–50. 10.1007/s13311-019-00816-2.

Carrera, Emmanuel, and Giulio Tononi. 2014. “Diaschisis: Past, Present, Future.” Brain 137 (9): 2408–22. 10.1093/brain/awu101.

Case, Theodore J. 1938. “ELECTROENCEPHALOGRAPHY IN THE DIAGNOSIS AND LOCALIZATION OF INTRACRANIAL CONDITIONS.” The Journal of Nervous and Mental Disease 87 (5): 598.

Cassidy, Jessica M., Anirudh Wodeyar, Jennifer Wu, Kiranjot Kaur, Ashley K. Masuda, Ramesh Srinivasan, and Steven C. Cramer. 2020. “Low-Frequency Oscillations Are a Biomarker of Injury and Recovery After Stroke.” Stroke 51 (5): 1442–50. 10.1161/STROKEAHA.120.028932.

Cuspineda, E., C. Machado, L. Galán, E. Aubert, M. A. Alvarez, F. Llopis, L. Portela, M. García, J. M. Manero, and Y. Ávila. 2007. “QEEG Prognostic Value in Acute Stroke.” Clinical EEG and Neuroscience 38 (3): 155–60. 10.1177/155005940703800312.

DeMarco, Andrew T., and Peter E. Turkeltaub. 2018. “A Multivariate Lesion Symptom Mapping Toolbox and Examination of Lesion-Volume Biases and Correction Methods in Lesion-Symptom Mapping.” Human Brain Mapping 39 (11): 4169–82. 10.1002/hbm.24289.

DeWitt, L. D., F. S. Buonanno, J. P. Kistler, T. J. Brady, I. L. Pykett, M. R. Goldman, and K. R. Davis. 1984. “Nuclear Magnetic Resonance Imaging in Evaluation of Clinical Stroke Syndromes.” Annals of Neurology 16 (5): 535–45. 10.1002/ana.410160503.

Drift, Jan Hendrik Andriaan van der. 1957. “The Significance of Electro-Encephalogarphy for the Dianose and Localisation of Cerebral Tumours.”

Evans, Alan C., Andrew L. Janke, D. Louis Collins, and Sylvain Baillet. 2012. “Brain Templates and Atlases.” NeuroImage, 20 YEARS OF fMRI, 62 (2): 911–22. 10.1016/j.neuroimage.2012.01.024.

Fanciullacci, Chiara, Federica Bertolucci, Giuseppe Lamola, Alessandro Panarese, Fiorenzo Artoni, Silvestro Micera, Bruno Rossi, and Carmelo Chisari. 2017. “Delta Power Is Higher and More Symmetrical in Ischemic Stroke Patients with Cortical Involvement.” Frontiers in Human Neuroscience 11. 10.3389/fnhum.2017.00385.

Flounders, Matthew W, Carlos González-García, Richard Hardstone, and Biyu J He. 2019. “Neural Dynamics of Visual Ambiguity Resolution by Perceptual Prior.” eLife 8 (March). 10.7554/eLife.41861.

Fox Michael D. 2018. “Mapping Symptoms to Brain Networks with the Human Connectome.” New England Journal of Medicine 379 (23): 2237–45. 10.1056/NEJMra1706158.

Gladstone, David J., Cynthia J. Danells, and Sandra E. Black. 2002. “The Fugl-Meyer Assessment of Motor Recovery after Stroke: A Critical Review of Its Measurement Properties.” Neurorehabilitation and Neural Repair 16 (3): 232–40. 10.1177/154596802401105171.

Griffis, Joseph C., Nicholas V. Metcalf, Maurizio Corbetta, and Gordon L. Shulman. 2021. “Lesion Quantification Toolkit: A MATLAB Software Tool for Estimating Grey Matter Damage and White Matter Disconnections in Patients with Focal Brain Lesions.” NeuroImage: Clinical 30 (January):102639. 10.1016/j.nicl.2021.102639.

Hebart, Martin N, Brett B Bankson, Assaf Harel, Chris I Baker, and Radoslaw M Cichy. 2018. “The Representational Dynamics of Task and Object Processing in Humans.” Edited by Jody C Culham. eLife 7 (January):e32816. 10.7554/eLife.32816.

Karnath, Hans-Otto, Christoph Sperber, and Christopher Rorden. 2019. “Reprint of: Mapping Human Brain Lesions and Their Functional Consequences.” NeuroImage 190 (April):4–13. 10.1016/j.neuroimage.2019.01.044.

Kaufmann, Brigitte C, Dario Cazzoli, Manuela Pastore-Wapp, Tim Vanbellingen, Tobias Pflugshaupt, Daniel Bauer, René M Müri, Tobias Nef, Paolo Bartolomeo, and Thomas Nyffeler. 2023. “Joint Impact on Attention, Alertness and Inhibition of Lesions at a Frontal White Matter Crossroad.” Brain 146 (4): 1467–82. 10.1093/brain/awac359.

Khanna, Preeya, and Jose M Carmena. 2015. “Neural Oscillations: Beta Band Activity across Motor Networks.” *Current Opinion in Neurobiology*, Large-Scale Recording Technology (32), 32 (June):60–67. 10.1016/j.conb.2014.11.010.

Knight, Robert T. 1984. “Decreased Response to Novel Stimuli after Prefrontal Lesions in Man.” Electroencephalography and Clinical Neurophysiology/Evoked Potentials Section 59 (1): 9–20. 10.1016/0168-5597(84)90016-9.

Knight, Robert T. 1997. “Distributed Cortical Network for Visual Attention.” Journal of Cognitive Neuroscience 9 (1): 75–91. 10.1162/jocn.1997.9.1.75.

Knight, Robert T., Steven A. Hillyard, David L. Woods, and Helen J. Neville. 1981. “The Effects of Frontal Cortex Lesions on Event-Related Potentials during Auditory Selective Attention.” Electroencephalography and Clinical Neurophysiology 52 (6): 571–82. 10.1016/0013-4694(81)91431-0.

Leyton, A. S. F., and C. S. Sherrington. 1917. “OBSERVATIONS ON THE EXCITABLE CORTEX OF THE CHIMPANZEE, ORANG-UTAN, AND GORILLA.” Quarterly Journal of Experimental Physiology 11 (2): 135–222. 10.1113/expphysiol.1917.sp000240.

Li, Hai, Gan Huang, Qiang Lin, Jiang-Li Zhao, Wai-Leung Ambrose Lo, Yu-Rong Mao, Ling Chen, Zhi-Guo Zhang, Dong-Feng Huang, and Le Li. 2018. “Combining Movement-Related Cortical Potentials and Event-Related Desynchronization to Study Movement Preparation and Execution.” Frontiers in Neurology 9. https://www.frontiersin.org/articles/10.3389/fneur.2018.00822.

Liew, Sook-Lei, Artemis Zavaliangos-Petropulu, Neda Jahanshad, Catherine E. Lang, Kathryn S. Hayward, Keith R. Lohse, Julia M. Juliano, et al. 2022. “The ENIGMA Stroke Recovery Working Group: Big Data Neuroimaging to Study Brain–Behavior Relationships after Stroke.” Human Brain Mapping 43 (1): 129–48. 10.1002/hbm.25015.

López-Larraz, Eduardo, Thiago C. Figueiredo, Ainhoa Insausti-Delgado, Ulf Ziemann, Niels Birbaumer, and Ander Ramos-Murguialday. 2018. “Event-Related Desynchronization during Movement Attempt and Execution in Severely Paralyzed Stroke Patients: An Artifact Removal Relevance Analysis.” NeuroImage: Clinical 20 (January):972–86. 10.1016/j.nicl.2018.09.035.

Lorca-Puls, Diego L., Andrea Gajardo-Vidal, Jitrachote White, Mohamed L. Seghier, Alexander P. Leff, David W. Green, Jenny T. Crinion, et al. 2018. “The Impact of Sample Size on the Reproducibility of Voxel-Based Lesion-Deficit Mappings.” Neuropsychologia 115 (July):101–11. 10.1016/j.neuropsychologia.2018.03.014.

Macdonell, R. A. L., G. A. Donnan, P. F. Bladin, S. F. Berkovic, and C. H. R. Wriedt. 1988. “The Electroencephalogram and Acute Ischemic Stroke: Distinguishing Cortical From Lacunar Infarction.” Archives of Neurology 45 (5): 520–24. 10.1001/archneur.1988.00520290048013.

Makin, Tamar R, and John W Krakauer. 2023. “Against Cortical Reorganisation.” Edited by J Andrew Pruszynski and Floris P de Lange. eLife 12 (November):e84716. 10.7554/eLife.84716.

Marek, Scott, Brenden Tervo-Clemmens, Finnegan J. Calabro, David F. Montez, Benjamin P. Kay, Alexander S. Hatoum, Meghan Rose Donohue, et al. 2022. “Reproducible Brain-Wide Association Studies Require Thousands of Individuals.” Nature 603 (7902): 654–60. 10.1038/s41586-022-04492-9.

Maris, Eric, and Robert Oostenveld. 2007. “Nonparametric Statistical Testing of EEG- and MEG-Data.” Journal of Neuroscience Methods 164 (1): 177–90. 10.1016/j.jneumeth.2007.03.024.

Mohr, J. P., W. C. Watters, and G. W. Duncan. 1975. “Thalamic Hemorrhage and Aphasia.” Brain and Language 2 (January):3–17. 10.1016/S0093-934X(75)80050-2.

Noble, Stephanie, Joshua Curtiss, Luiz Pessoa, and Dustin Scheinost. 2023. “The Tip of the Iceberg: A Call to Embrace Anti-Localizationism in Human Neuroscience Research.” PsyArXiv. 10.31234/osf.io/9eqh6.

Oostenveld, Robert, Pascal Fries, Eric Maris, and Jan-Mathijs Schoffelen. 2010. “FieldTrip: Open Source Software for Advanced Analysis of MEG, EEG, and Invasive Electrophysiological Data.” Computational Intelligence and Neuroscience 2011 (December):e156869. 10.1155/2011/156869.

Penfield, Wilder, and Theodore Rasmussen. 1950. The Cerebral Cortex of Man; a Clinical Study of Localization of Function. The Cerebral Cortex of Man; a Clinical Study of Localization of Function. Oxford, England: Macmillan.

Pernet, Cyril R., Stefan Appelhoff, Krzysztof J. Gorgolewski, Guillaume Flandin, Christophe Phillips, Arnaud Delorme, and Robert Oostenveld. 2019. “EEG-BIDS, an Extension to the Brain Imaging Data Structure for Electroencephalography.” Scientific Data 6 (1): 103. 10.1038/s41597-019-0104-8.

Pustina, Dorian, Brian Avants, Olufunsho K. Faseyitan, John D. Medaglia, and H. Branch Coslett. 2018. “Improved Accuracy of Lesion to Symptom Mapping with Multivariate Sparse Canonical Correlations.” Neuropsychologia 115 (July):154–66. 10.1016/j.neuropsychologia.2017.08.027.

Ramanathan, Dhakshin S., Ling Guo, Tanuj Gulati, Gray Davidson, April K. Hishinuma, Seok-Joon Won, Robert T. Knight, Edward F. Chang, Raymond A. Swanson, and Karunesh Ganguly. 2018. “Low-Frequency Cortical Activity Is a Neuromodulatory Target That Tracks Recovery after Stroke.” Nature Medicine 24 (8): 1257–67. 10.1038/s41591-018-0058-y.

Raposo, Isabel, Sara M. Szczepanski, Kathleen Haaland, Tor Endestad, Anne-Kristin Solbakk, Robert T. Knight, and Randolph F. Helfrich. 2023. “Periodic Attention Deficits after Frontoparietal Lesions Provide Causal Evidence for Rhythmic Attentional Sampling.” *Current Biology*, October. 10.1016/j.cub.2023.09.065.

Ray, Andreas M., Eduardo Lopez-Larraz, Thiago C. Figueiredo, Niels Birbaumer, and Ander Ramos- Murguialday. 2017. “Movement-Related Brain Oscillations Vary with Lesion Location in Severely Paralyzed Chronic Stroke Patients.” In 2017 39th Annual International Conference of the IEEE Engineering in Medicine and Biology Society (EMBC), 1664–67. Seogwipo: IEEE. 10.1109/EMBC.2017.8037160.

Rogers, Jeffrey, Sandy Middleton, Peter H. Wilson, and Stuart J. Johnstone. 2020. “Predicting Functional Outcomes after Stroke: An Observational Study of Acute Single-Channel EEG.” Topics in Stroke Rehabilitation 27 (3): 161–72. 10.1080/10749357.2019.1673576.

Rossiter, Holly E., Marie-Hélène Boudrias, and Nick S. Ward. 2014. “Do Movement-Related Beta Oscillations Change after Stroke?” Journal of Neurophysiology 112 (9): 2053–58. 10.1152/jn.00345.2014.

Saes, Mique, Sarah B. Zandvliet, Aukje S. Andringa, Andreas Daffertshofer, Jos W. R. Twisk, Carel G. M. Meskers, Erwin E. H. van Wegen, and Gert Kwakkel. 2020. “Is Resting-State EEG Longitudinally Associated With Recovery of Clinical Neurological Impairments Early Poststroke? A Prospective Cohort Study.” Neurorehabilitation and Neural Repair 34 (5): 389–402. 10.1177/1545968320905797.

Sharp, Frank R., Aigang Lu, Yang Tang, and David E. Millhorn. 2000. “Multiple Molecular Penumbras after Focal Cerebral Ischemia.” Journal of Cerebral Blood Flow & Metabolism 20 (7): 1011–32. 10.1097/00004647-200007000-00001.

Shreve, Lauren, Arshdeep Kaur, Christopher Vo, Jennifer Wu, Jessica M. Cassidy, Andrew Nguyen, Robert J. Zhou, et al. 2019. “Electroencephalography Measures Are Useful for Identifying Large Acute Ischemic Stroke in the Emergency Department.” Journal of Stroke and Cerebrovascular Diseases: The Official Journal of National Stroke Association 28 (8): 2280–86. 10.1016/j.jstrokecerebrovasdis.2019.05.019.

Smith, Stephen M., and Thomas E. Nichols. 2009. “Threshold-Free Cluster Enhancement: Addressing Problems of Smoothing, Threshold Dependence and Localisation in Cluster Inference.” NeuroImage 44 (1): 83–98. 10.1016/j.neuroimage.2008.03.061.

Spisak, Tamas, Ulrike Bingel, and Tor D. Wager. 2023. “Multivariate BWAS Can Be Replicable with Moderate Sample Sizes.” Nature 615 (7951): E4–7. 10.1038/s41586-023-05745-x.

Taha, Aladdin, Joaquim Bobi, Ruben Dammers, Rick M. Dijkhuizen, Antje Y. Dreyer, Adriaan C.G.M. van Es, Fabienne Ferrara, et al. 2022. “Comparison of Large Animal Models for Acute Ischemic Stroke: Which Model to Use?” Stroke 53 (4): 1411–22. 10.1161/STROKEAHA.121.036050.

Tang, Chih-Wei, Fu-Jung Hsiao, Po-Lei Lee, Yun-An Tsai, Ya-Fang Hsu, Wei-Ta Chen, Yung-Yang Lin, Charlotte J. Stagg, and I-Hui Lee. 2020. “β-Oscillations Reflect Recovery of the Paretic Upper Limb in Subacute Stroke.” Neurorehabilitation and Neural Repair 34 (5): 450–62. 10.1177/1545968320913502.

Thiebaut de Schotten, Michel, Chris Foulon, and Parashkev Nachev. 2020. “Brain Disconnections Link Structural Connectivity with Function and Behaviour.” Nature Communications 11 (1): 5094. 10.1038/s41467-020-18920-9.

Van Der Drift, J.H.A. 1961. “Ischemic Cerebral Lesions.” Angiology 12 (9): 401–18. 10.1177/000331976101200902.

Voytek, Bradley, Matar Davis, Elena Yago, Francisco Barceló, Edward K. Vogel, and Robert T. Knight. 2010. “Dynamic Neuroplasticity after Human Prefrontal Cortex Damage.” Neuron 68 (3): 401–8. 10.1016/j.neuron.2010.09.018.

Walter, W. Grey. 1936. “THE LOCATION OF CEREBRAL TUMOURS BY ELECTRO-ENCEPHALOGRAPHY.” *The Lancet*, Originally published as Volume 2, Issue 5893, 228 (5893): 305 –8. 10.1016/S0140-6736(01)05173-X.

Ward, Nick S., Jennifer M. Newton, Orlando B. C. Swayne, Lucy Lee, Richard S. J. Frackowiak, Alan J. Thompson, Richard J. Greenwood, and John C. Rothwell. 2007. “The Relationship between Brain Activity and Peak Grip Force Is Modulated by Corticospinal System Integrity after Subcortical Stroke.” The European Journal of Neuroscience 25 (6): 1865–73. 10.1111/j.1460-9568.2007.05434.x.

Wernicke, Carl. 1874. Der aphasische Symptomencomplex: eine psychologische Studie auf anatomischer Basis. Cohn & Weigert.

Wiesen, Daniel, Hans-Otto Karnath, and Christoph Sperber. 2020. “Disconnection Somewhere down the Line: Multivariate Lesion-Symptom Mapping of the Line Bisection Error.” Cortex; a Journal Devoted to the Study of the Nervous System and Behavior 133 (December):120–32. 10.1016/j.cortex.2020.09.012.

Williams, Denis, and Frederic A. Gibbs. 1938. “The Localization of Intracranial Lesions by Electroencephalography.” New England Journal of Medicine 218 (24): 998–1002. 10.1056/NEJM193806162182402.

Wu, Jennifer, Erin Burke Quinlan, Lucy Dodakian, Alison McKenzie, Nikhita Kathuria, Robert J. Zhou, Renee Augsburger, et al. 2015. “Connectivity Measures Are Robust Biomarkers of Cortical Function and Plasticity after Stroke.” Brain 138 (8): 2359–69. 10.1093/brain/awv156.

Wu, Jennifer, Ramesh Srinivasan, Erin Burke Quinlan, Ana Solodkin, Steven L. Small, and Steven C. Cramer. 2016. “Utility of EEG Measures of Brain Function in Patients with Acute Stroke.” Journal of Neurophysiology 115 (5): 2399–2405. 10.1152/jn.00978.2015.

Zhang, Yongsheng, Daniel Y. Kimberg, H. Branch Coslett, Myrna F. Schwartz, and Ze Wang. 2014. “Multivariate Lesion-Symptom Mapping Using Support Vector Regression.” Human Brain Mapping 35 (12): 5861–76. 10.1002/hbm.22590.

Ziegler, Dewey K. 1954. “Cerebral Atrophy in Psychiatric Patients.” American Journal of Psychiatry 111 (6): 454–58. 10.1176/ajp.111.6.454.

